# Radiopotentiation of Enzalutamide Over Human Prostate Cancer Cells as Assessed by Real-Time Cell Monitoring

**DOI:** 10.1101/324889

**Authors:** Marta Barrado, Idoia Blanco-Luquin, Navarrete Paola Andrea, Ignacio Visus, David Guerrero-Setas, David Escors, Grazyna Kochan, Fernando Arias

**Affiliations:** Biomedical Research Center of Navarra-Navarrabiomed. Fundación Miguel Servet. IdlSNA. Irunlarrea 3, 31008 Pamplona, Navarre, Spain.; Department of Radiation Oncology. Hospital of Navarre. IdISNA. Irunlarrea 3, 31008 Pamplona, Navarre, Spain.; Department of Infection and Immunity, Rayne Insitute, University College London, 5 University Street WC1E 6JJ London, United Kingdom.

**Keywords:** Enzalutamide, radiotherapy, apoptosis, prostate cancer, androgen blockade

## Abstract

While radiotherapy is the first line of treatment for prostate cancer, androgen blockade therapies are demonstrating significant survival benefit as monotherapies. As androgen blockade can cause cell death by apoptosis, it is likely that androgen blockade will potentiate the cytotoxic activities of radiotherapy. Here we tested the potential synergistic effects of these two treatments over two human metastatic prostate cancer cells by real time growth monitoring (RTCA), androgen-sensitive LNCaP cells and androgen-resistant PC-3. Both cell lines were highly resistant to high doses of radiotherapy. A pre-treatment of LNCaP cells with IC50 concentrations of enzalutamide significantly sensitized them to radiotherapy through enhanced apoptosis. In contrast, enzalutamide resistant PC-3 cells were not sensitized to radiotherapy by androgen blockade. These results provide evidence that the enzalutamide/radiotherapy combination could maximize therapeutic responses in patients with enzalutamide-sensitive prostate cancer.

## INTRODUCTION

Radiotherapy is the most widely used first-line treatment for prostate cancer. Approximately 50% of the patients are treated with radiotherapy alone or in combination with other therapies. However, a significant percentage of patients relapse ^1^. Androgen receptor (AR)-targeted therapies have been developed to prevent proliferation and induce apoptosis in prostate cancer cells ^2^, and these are demonstrating significant survival benefit as monotherapies. This is certainly the case of enzalutamide, a highly selective AR antagonist that blocks its association with androgen, preventing its nuclear translocation and inhibiting its role as a transcription factor. Nevertheless, primary and secondary resistances eventually develop through a series of mechanisms that include AR mutations that hamper binding to enzalutamide ^3^. However, preclinical findings suggest that its combination with traditional treatments may achieve additive or synergistic effects. These effects should most likely reduce the appearance of resistance to AR blockade and augment patient benefit ^4^ Hence, a collection of combinatorial treatments with enzalutamide are currently being tested such as mTOR and HER2 inhibition ^5^ and antagonists of apoptotic inhibitors amongst other agents ^6^.

Radiotherapy is an efficacious inducer of cell death and enzalutamide treatment should further sensitize irradiated prostate cancer cells to induction of apoptosis. However, this combination has not been tested yet in clinical practice as there is a lack of preclinical data validating this approach.

Most of the studies on apoptotic agents employ end-point MTT cytotoxicity assays or clonogenic assays which show multiple drawbacks. Hence, impedance-based technologies have been developed to monitor cell growth/viability in real time, such as the xCelligence RTCA technology ^7^ This technology possesses the sensibility to detect changes in cell size, shape and growth, making it a very useful technique for monitoring effects of pharmacological agents on cellular biology^7,8^ Moreover, RTCA integrates the real-time data to plot inhibition curves to accurately calculate inhibitory doses (ID) at any given time. This is in contrast to MTT assays that rely on end-points to calculate cell viability, or clonogenic assays that are difficult to interpret. In this study we have addressed the potential of RTCA technology to test whether radiotherapy/enzalutamide combinatorial regimes in prostate cancer cell lines may warrant their application in human patients. Our data provides the basis for using enzalutamide as an agent to potentiate radiotherapy for prostate cancer.

## RESULTS

### Estimation of enzalutamide IC50s on two human metastatic prostate cancer cell lines by real-time cell monitoring

We analyzed the sensitivity to enzalutamide of two human prostate cancer cell lines extensively used in preclinical work by real-time monitoring of cell growth (xCelligence RTCA). The LNCaP cell line was derived from a lymph node metastasis of a 50-year old male Caucasian patient. These cells are androgen and estrogen receptor-positive and also sensitive to androgen blockade. PC-3 cells were chosen as androgen insensitive control cells which were obtained from a bone metastasis of a 62-year old male Caucasian patient with a grade IV prostate adenocarcinoma.

First, we validated the use of real-time cell monitoring to estimate the sensitivity of the two cell lines to enzalutamide, as most of the studies utilize end-point-based assays. Concentrations of enzalutamide ranging from 0 to 10 μM were added to cell cultures and their growth was monitored up to 12 days. DMSO was added as a control and treatments were carried out in duplicates and repeated twice. The growth of LNCaP cells exhibited exponential kinetics which was highly significantly delayed with increasing enzalutamide concentrations, especially from 5 μM onwards (**Fig. 1a**). In contrast, no significant effects were observed in the growth of PC-3 cells at any of the tested enzalutamide concentrations, in agreement with their intrinsic resistance to enzalutamide (Fig. 1b). Then, IC50 concentrations were estimated with RTCA data for the two cell lines (Fig. 1c). The calculated IC50 for LNCaP cells (5.6±0.8 μM; n=4) was comparable to that published by other authors using end-point cytotoxicity assays (between 1 and 5 μM) ^9,10^. In contrast, the calculated IC50 value for PC-3 cells had to be extrapolated up to a non-physiological value (34.9±9 μM; n=4). This was consistent with the intrinsic resistance of PC-3 cells to androgen blockade (Fig. 1c).

**Figure 1.**
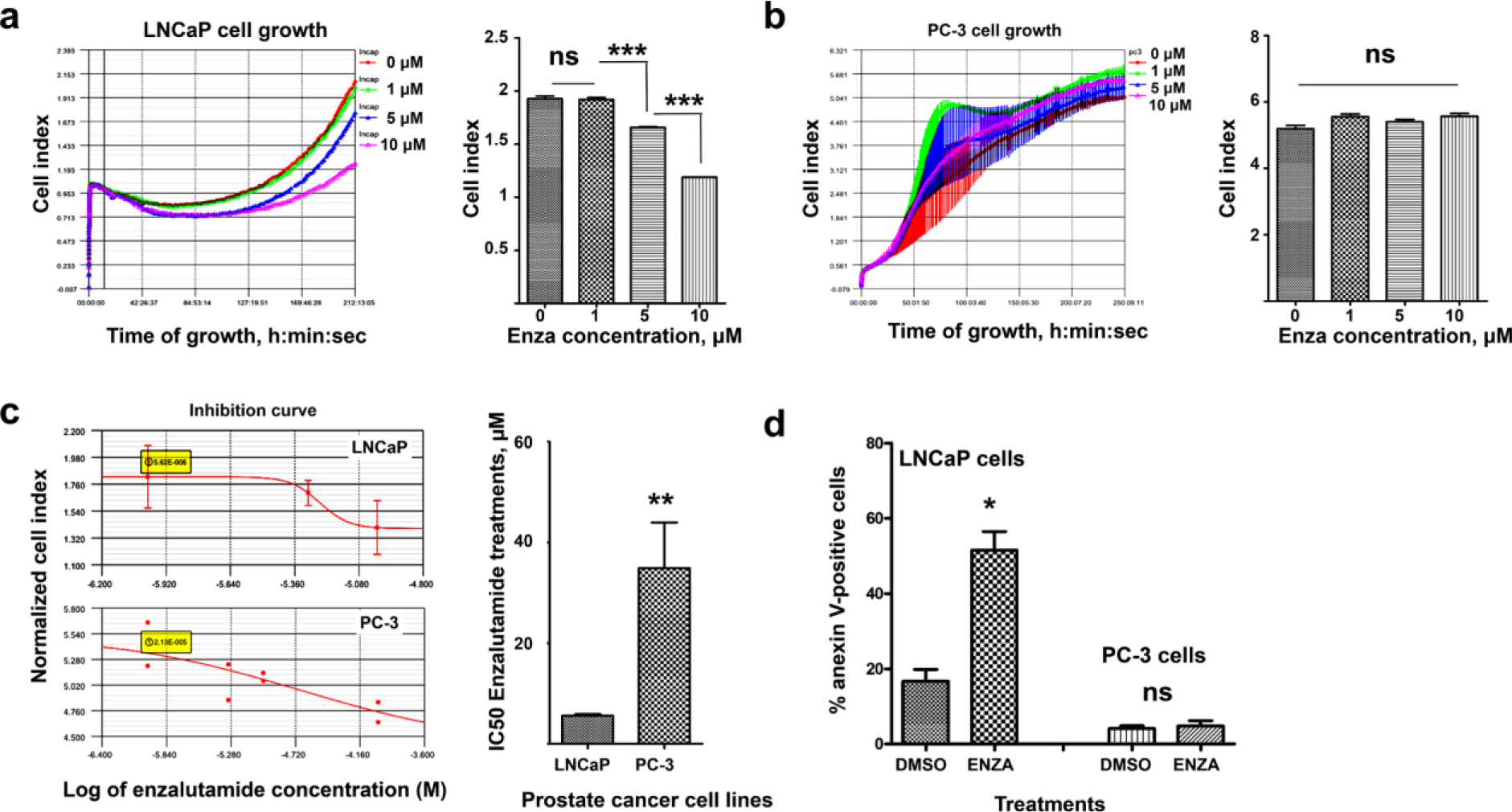
Inhibition of cell growth by enzalutide as assessed by real-time cell monitoring. (**a**) The graph on the left shows real-time growth kinetics from control-treated LNCaP cells (DMSO carrier) or treated with 1, 5 or 10 μM enzalutamide, as indicated in the graph. On the right, the data from two RTCA experiments with duplicates within each one are plotted as a bar graph, with means (Cell Indexes) and standard deviations as error bars. (**b**) As in (a) but with PC-3 cells. Cell index is plotted as means from duplicates together with error bar. (**c**) The graphs on the left show the inhibition plots of either LNCaP cells (top graph) or PC-3 cells (bottom graph) following enzalutamide treatments at increasing concentrations, as calculated by RTCA. The calculated IC50 values are highlighted in yellow within each graph. On the right, column graphs representing the calculated IC50s from two independent experiments, each experiment in duplicates, for LNCaP and PC-3 cells as indicated. Relevant statistical comparisons are shown. (**d**) Bar graph representing the percentage of anexin V-positive cells in cultures of the indicated cell lines, treated with DMSO or enzalutamide at the IC50 concentrations as calculated in (c). Relevant statistical comparisons are shown within the graphs. *, **, ***, indicate significant (P<0.05), very significant (P<0.01) and highly (P<0.001) significant differences, respectively.

Real-time cell monitoring is based on impedance readings from cell culture plates on which cells attach and proliferate^7^. Although it can be used to test cell viability, this technique does not directly provide information on cell death. Therefore, we tested the induction of apoptosis by enzalutamide in prostate cancer cells by anexin V staining followed by flow cytometry. Hence, the two cell lines were treated with DMSO or enzalutamide for 48 hours at their corresponding IC50 concentrations (Fig. 1c). Then, the percentage of anexin V-positive cells was quantified by flow cytometry, and it was confirmed that enzalutamide significantly increased apoptosis in treated LNCaP cells (Fig. 1D). As expected, PC3 cells were refractory to apoptosis induction by enzalutamide.

### Enzalutamide pre-treatment potentiates radiotherapy

Prostate cancer cell lines were extraordinarily resistant to radiotherapy compared to other non-prostate cancer cell lines (data not shown). Therefore, we tested whether the pro-apoptotic effects of enzalutamide could also sensitize cells to radiotherapy-induced cell death. To test this, LNCaP and PC-3 cells were pre-treated with DMSO or enzalutamide at their IC50 concentrations for 3 days. At these concentrations, cell growth is delayed but cells remain mostly viable. Then, cells were irradiated with 10, 20 and 30 Gy and their growth and viability were monitored by RTCA. As expected from our preliminary studies, doses of 30 Gy had to be used to induce measurable cell death in untreated LNCaP cells (Fig. 2a). In contrast, pretreatment of LNCaP cells with enzalutamide strongly sensitized LNCaP cells to irradiation causing rapid cell death at the minimum tested dose of 10 Gy. Enzalutamide-resistant PC-3 exhibited exponential growth kinetics at the three tested irradiation doses, and pre-treatment with 40 μM enzalutamide did not have any effect whatsoever to their growth (Fig. 2b). These results strongly indicated that enzalutamide sensitized prostate cancer cells to radiotherapy as long as they respond to androgen blockade. To evaluate if these synergistic effects were caused by increased apoptosis, anexin V staining followed by flow cytometry was used. As observed by RTCA, LNCaP cells pre-treated by DMSO were highly resistant to radiotherapy-induced apoptosis at a dose of 10 Gy. In contrast, pre-treatment with ID50 concentrations of enzalutamide very highly sensitized cells to radiotherapy-induced apoptosis at a dose of 10 Gy (Fig. 2c). PC-3 cells remained insensitive to enzalutamide, radiotherapy or their combination, suggesting that enzalutamide′s apoptotic properties were required for radiopotentiation.

**Figure 2.**
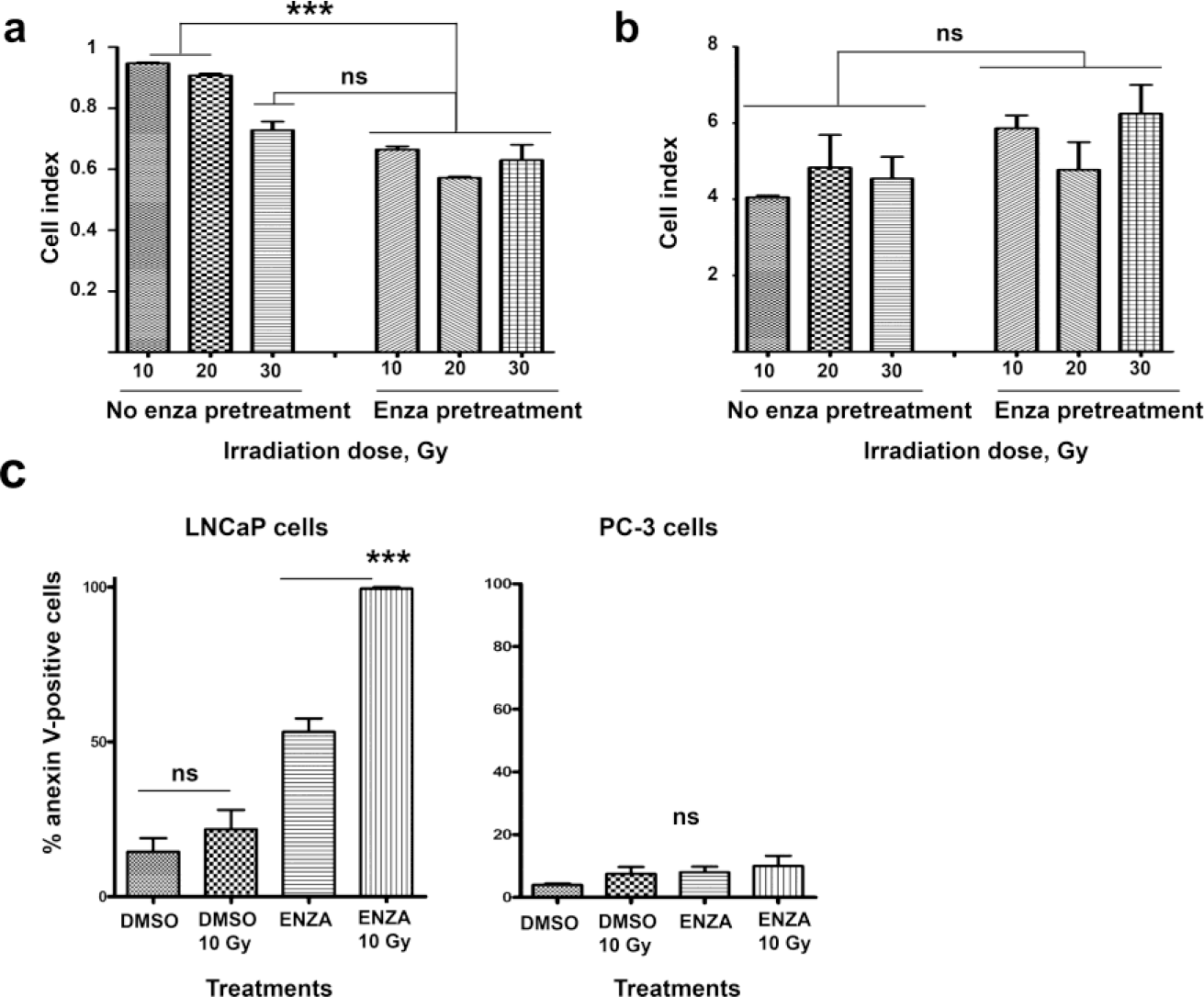
Enzalutamide sensitizes LNCaP cells but not PC-3 cells to radiotherapy. (**a**) RTCA cell growth data of LNCaP cells treated with either DMSO (vehicle control, graph on the left) or enzalutamide (graph on the right) for three days followed by the irradiation doses as indicated. The bar graph represents means of Cell Index with standard deviations as error bars. Treatments with DMSO (no enza) or enzalutamide are indicated, as well as irradiation doses. (**b**) Same as in (a) but for PC-3 cells. (**c**) Bar graph representing the percentage of anexin V-positive cells in cultures of the indicated cell lines, treated with DMSO or enzalutamide at the IC50 concentrations followed by treatments with 10 Gy where indicated. Relevant statistical comparisons are shown within the graphs. ***, indicates highly significant differences (P<0.001). ns, indicates non-significant differences (P>0.05).

## DISCUSSION

Prostate cancer therapy heavily relies on androgen deprivation which has improved survival ^11^. Nevertheless, disease progression still occurs (castration-resistant prostate cancer), although there is evidence that in this case cancer progression continues to be driven by androgen signaling ^11^. To reinforce the suppression of AR signaling, several AR antagonists have been developed. Amongst these, enzalumtamide (MDV3100) is an inhibitor that binds to the ligand-binding domain of AR and blocks its nuclear translocation ^12^ Enzalutamide shows also benefit for the treatment of castration-resistant prostate cancer ^13^.

As radiotherapy is the most widespread first line treatment for prostate cancer, its combination with enzalutamide represents a rather logical choice. However, clinical studies addressing this combination are scarce due to lack of pre-clinical evidence on the benefits of this combination. In this study we have applied impedance-based real-time cell monitoring to evaluate the potential of the enzalutamide/radiotherapy combination over two human prostate cancer cell lines; Enzalutamide sensitive LNCaP cells and enzalutamide-resistant PC-3 cells. These two cell lines are extensively used in pre-clinical evaluation of prostate cancer treatments. Our RTCA data confirmed the IC50s of enzalutamide for LNCaP cells estimated by other authors using end-point based assays, as well as the enzalutamide-resistance of PC-3. Enzalutamide increased apoptosis in LNCaP cell cultures, and suggested that this property may further sensitize these cells to radiotherapy. In fact, we found that LNCaP and PC-3 cells were strongly resistant to radiotherapy. Indeed, PC-3 cells remained viable even at the highest irradiation dose tested in this study (30 Gy), while LNCaP cells only died at this very high dose.

Interestingly, a short pre-treatment of LNCaP cells with enzalutamide at the IC50 concentration strongly sensitized them to radiotherapy, which induced cell death by apoptosis at the lower dose tested in this study (10 Gy). In contrast PC-3 cells remained insensitive to irradiation in the presence of enzalutamide. However, these results strongly suggested that the pro-apoptotic properties of enzalutamide were absolutely required for its radiopotentiating effects, only over androgen blockade-sensitive cells. Additionally, we demonstrate that RTCA is a simple and straightforward method to evaluate cell death by radiotherapy. This technique has major advantages compared to MTT or clonogenic assays. These assays show high variability and in many instances lack of reproducibility.

Based on our findings, the combination of radiotherapy with enzalutamide represents a good basis for ulterior clinical trials in locally advanced prostate cancer. Moreover, the irradiation doses could be lowered for patients with enzalutamide-sensitive tumors.

## METHODS

### Cells

LNCaP and PC3 human prostate cancer cell lines were purchased from the ATCC and grown as recommended by the manufacturer. LNCaP cells were grown in RPMI-1640 medium. PC-3 cells were grown in F-12 medium. All culture media were supplemented with 10% FCS, penicillin and streptomycin.

### Radiotherapy

Triplicate prostate cancer cell cultures were subjected to 3 different doses of radiotherapy, 10, 20 and 30 Gy in the Radiation Oncology Department of the Hospital of Navarre in collaboration with the Physics department.

### Flow cytometry

Cells were collected and stained using the eBioscience Anexin V apoptosis assay as described by the manufacturer. Data was collected with a FACS CANTO (BD bioscience). Experiments were performed in duplicates.

### Real-Time Cell Monitoring (RTCA)

Cell growth and viability were monitored by XCelligence RTCA (ACEA) as described before ^14^ for up to two weeks of culture. Inhibition curves were obtained using the in-built ACEA software with RTCA data.

### Statistics

RTCA cultures were carried out in duplicates and experiments repeated twice. The Cell Index data was highly reproducible and normally distributed. Therefore, the treatment groups were compared with one-way ANOVA using the means from duplicates and data from all replicate experiments, during the last 24 h of culture. Pair-wise comparisons were carried out with Tukey’s test. Data on Anexin-V staining was compared using the non-parametric test of Mann-Whitney, as the variances of the groups were significantly different. GraphPad Prism was used to carry out statistical tests.

### Data Availability Statement

Reagent or cell line derived from this study will be provided when requested.

## ACKNOWLEDGEMENTS

This project was funded by a research project grant from Astellas Oncology Inc., and partially by a FIS grant (PI14/00579) from the Instituto de Salud Carlos III, a grant from the Government of Navarre (BMED 033-2014, Spain) and a grant from Fundación Caixa (Spain) to GK. We thank SARAY foundatión (Navarra, Spain) and Sandra Ibarra′s foundation for their financial support. DE is funded by a Miguel Servet Fellowship (CP12/03114) awarded by the Instituto de Salud Carlos III (Spain).

## AUTHOR CONTRIBUTIONS

MB, IBL, PAN, IV and DE performed experiments and analyzed data. DGS, DE, GK and FA designed the study, performed experiments and analyzed the data. All authors contributed to the writing of the manuscript.

